# Accurate 3D SMLM localization via Vectorial In-situ PSF Retrieval and Aberration Assessment

**DOI:** 10.1101/2023.11.03.565592

**Authors:** Xinxun Yang, Hongfei Zhu, Yile Sun, Hanmeng Wu, Yubing Han, Xiang Hao, Renjie Zhou, Cuifang Kuang, Xu Liu

## Abstract

In single-molecule localization microscopy (SMLM), achieving precise localization hinges on obtaining an authentic point spread function (PSF) influenced by system and sample-induced aberrations. Here, we introduce VISPR (Vectorial in-situ PSF retrieval) retrieving precise 3D PSF models considering both system and sample-induced aberrations under SMLM conditions. By employing the theory of vectorial PSF model and maximum likelihood estimation (MLE) phase retrieval, VISPR is capable of reconstructing an accurate 3D PSF model achieving the theoretically minimum uncertainty and accurately reflecting three-dimensional information of single molecules. This capability empowers accurate 3D super-resolution reconstruction in 3D SMLM. Additionally, VISPR applies to low signal-to-noise ratio circumstances and is adept at retrieving high-frequency details of the experimental PSF across an extensive depth range—a challenging feat for alternative approaches. As an effective tool, VISPR enables the quantitative assessment of aberrations induced by the system and sample environment. From the simulations and experiments, we verified the superiority and effectiveness of VISPR. It is essential to highlight that VISPR applies to various SMLM microscope modalities.

## 1. Introduction

In recent decades, super-resolution fluorescent microscopy techniques have become a crucial platform that surmounted the diffraction limit for exploring cellular structures, including proteins, lipids, DNA, and RNA at the nanoscale. These technologies have played an important role in the development of cell biology science. To be specific, Single Molecule Localization Microscopy (SMLM[1–3]), exemplified by techniques like PALM[2]/STORM[3], employs photo-switchable or blinking dyes and proteins to precisely detect and localize individual molecules within a sub-diffraction region, achieving sub-5nm precision in three dimensions[4, 5]. In 3D SMLM[6–12], the critical factor is the accurate knowledge of the Point Spread Function (PSF), enabling the precise localization and description of molecules with respect to their axial position within the specimen for 3D super-resolution imaging[13–15]. Besides, acquiring an accurate PSF holds significant importance for multiple applications, including the correction and analysis of aberrations introduced by the instrument and sample environment[16, 17], quantification and enhancement of microscopy system performance[18, 19], point spread functions (PSFs) engineering, such as double helix[8], phase ramp[20], tetrapod[21] or Biplane configuration[12], and the enhancement of SMLM localization precision. Hence, it is crucial to attain an accurate PSF model that accounts for the impact of instrument imperfections and sample-induced aberrations arising from the non-uniform refractive indices within the sample.

To obtain the PSF model, current approaches rely on the calibrations generated from fiducial markers such as beads or gold nanoparticles[16, 22–27], which takes bead stacks moving at certain z-axis intervals and subsequently utilizing phase retrieval algorithms for PSF retrieval. However, photons emitted by these fiducial markers never traverse the cellular or tissue sample, thus failing to consider aberrations induced by the sample environment, such as refraction mismatch caused by the photons passing through the complex biological and optical environment. Besides, the bead size ranges from 40 to 200 nm, significantly exceeding the size of a fluorophore molecule, which is approximately 2 nm. Therefore, the PSF model retrieved by beads has limitations when localizing in SMLM, such as heightened uncertainty in localization accuracy and resulting positional deviations in 3D.

Recently, Xu *et al.* introduced INSPR generating in-situ PSF model directly derived from raw data in 3D SMLM using a modified Gerchberg-Saxton (GS) algorithm[28], which takes into account both the system-induced and sample-induced aberrations and is well-suited to the SMLM experimental conditions. However, a significant limitation of Xu’s algorithm lies in its use of a scalar PSF model, rendering it unsuitable for high numerical aperture (NA) microscope systems and leading to model deviations during PSF generation. Additionally, the GS algorithm is susceptible to noise, making it unsuitable for data with a low signal-to-noise ratio[29]. Moreover, INSPR incorporates an empirical Optical Transfer Function (OTF) scaling function in data processing[30], resulting in the exclusion of high-frequency information within the PSF spatial domain. This omission causes the generated PSF to gradually deviate from the authentic PSF as the depth axis increases. Furthermore, this rescaling strategy lacks consistency in phase restoration and cannot be theoretically supported or analyzed[29]. Consequently, the accuracy of INSPR is constrained by model deviation, the suppression of PSF high-frequency information, and sensitivity to background noise.

To accurately retrieve all parameters of a PSF model and address the issues mentioned above, we draw inspiration from Xu’s approach[28] and introduce the VISPR (Vectorial in-situ PSF retrieval), which is founded on a vectorial PSF model and Maximum Likelihood Estimation (MLE) phase estimation. In the article, we first introduce the theoretical foundations of VISPR, including the introduction of the vector PSF model and the MLE algorithm. Subsequently, we employ both simulation experiments and practical experiments to validate the efficacy of VISPR. VISPR effectively addresses the issues in Xu’s method, and accurately retrieves the 3D PSF model, achieving the theoretical Cramér-Rao lower bound (CRLB) and faithfully reflecting three-dimensional information of single molecules. This enhancement leads to improved accuracy in the reconstruction of 3D SMLM. Furthermore, our methodology exhibits robustness under low signal-to-noise ratio circumstances and proves adaptable to a diverse range of systems capable of SMLM experiments.

## 2. Principles and Methods

### 2.1. Microscope setup

To implement and validate VISPR, we built a customized SMLM system based on adaptive optics as shown in Figure S1. By going through an optical coupler (Schäfter + Kirchhoff GmbH, 60SMS-1-4-RGBV11-47), the single-wavelength Laser was coupled into a single-mode optical fiber (Schäfter + Kirchhoff GmbH, PMC-E-400RGB-3.5-NA011-3-APC.EC-350-P) and then collimated by a beam collimator (Schäfter + Kirchhoff GmbH, 60FC-T-4-M25-01). after the module of fiber optics, the beam propagated to a 200-mm tube lens (Thorlabs, TTL200-A) with relay of a 4f system which adjusted the illumination area and made the pixel size to be 89 nm. Through the tube lens, the excitation beam was focused to the back focal plane of an oil-immersion objective lens (Olympus, UPLXAPO100XO).) and formed epi-illumination. The fluorescence of each single molecule was collected by objective lens and passed to a deformable mirror (Boston Micromachines Corporation, Multi-3.5L-DM). It should be noted that the surface of deformable mirror conjugated with back focal plane and can be relayed by 4f configuration. Finally, the pupil was focused by a doublet lens (Thorlabs, AC254-125-A-ML) and the single-molecule raw images were collected by a scientific CMOS (Hamamatsu, ORCA-FusionBT C15440-20UP).

### 2.2. VISPR Principle

VISPR overcomes the limitations inherent in phase retrieval methods based on beads[16, 22–27] by directly retrieving the 3D PSF model from the raw single-molecule blinking dataset obtained from 3D SMLM experiments. The fundamental concept of 3D-SMILE is illustrated in Figure. 1. It consists of three significant steps: 1) PSF library construction, wherein the SMLM dataset is segmented into a single-spot PSF library representing a random axial distribution of the 3D PSF to be retrieved; 2) PSF library assignment, involving the utilization of an ideal vectorial PSF (generated by a constant pupil and using a vectorial PSF forward model) as a reference PSF to categorize the PSF library into classes based on axial information, employing mathematical frameworks of expectation-maximization[31] and *k*-means[32] to calculate the similarity between images; 3) 3D PSF model estimation, wherein temporarily axially assigned single-molecule spots are assembled, aligned, and averaged to form a 3D PSF stack. Subsequently, Maximum Likelihood Estimation (MLE) phase retrieval is employed to estimate the new pupil and Zernike coefficients of the PSF stack. This updated pupil generates a refined vectorial reference template, and steps 2-3 are iteratively repeated. For data with low signal-to-noise ratio (SNR), 6-10 iterations are required to converge. To avoid the PSF degeneracies that occurred during the process mentioned in Xu[28], we use the prior knowledge of the astigmatism orientation to generate asymmetric reference PSF, which helps us to build the unique in-situ vectorial PSF model.

**Figure 1.**
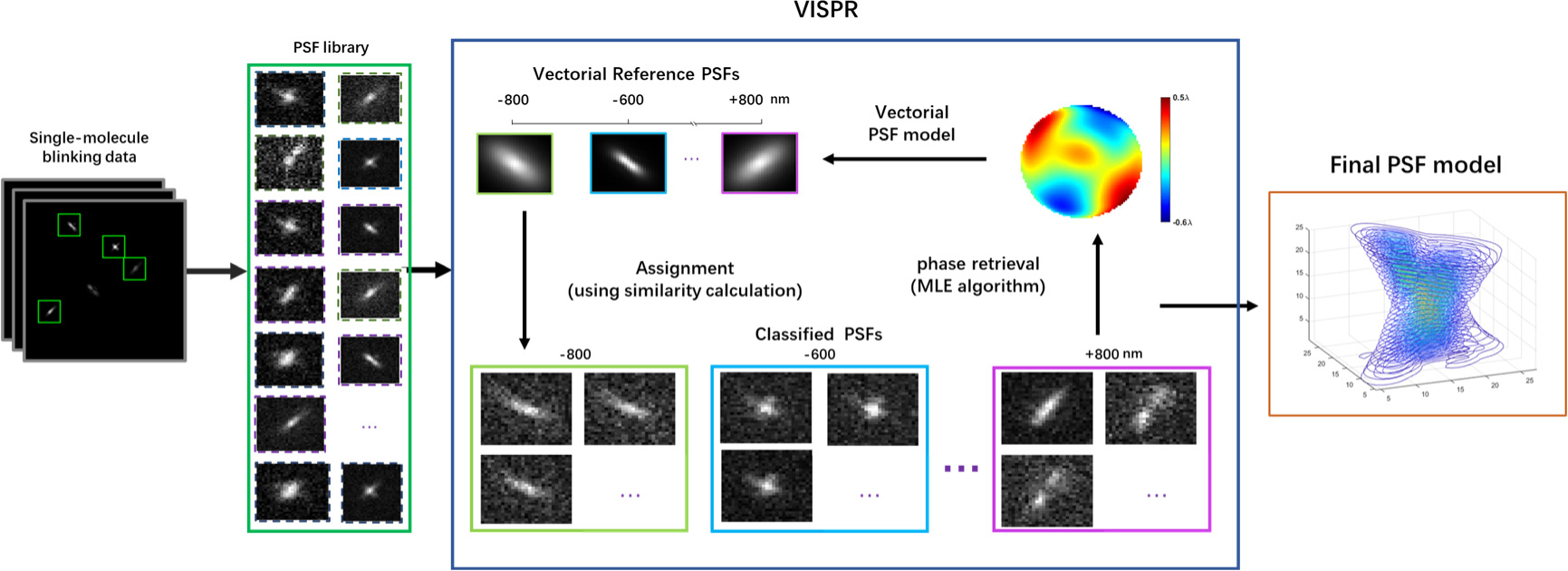
Concept of VISPR. After the single-molecule dataset (left) is acquired in SMLM experiments, a PSF library is obtained by segmentation. VISPR uses a starting pupil function with constant Zernike coefficients to obtain the reference vector PSF stacks. Then the reference PSF stacks assigns each emitter pattern to different z-axis positions according to its similarity with the template. These axially assigned PSFs are subsequently grouped, aligned and averaged to form a 3D PSF stack, which is then used to estimate the PSF model using MLE phase retrieval. The new pupil generates an updated reference vectorial PSF model for the next iteration. This process iterates about 6-8 times to converge and get the final PSF model.

#### 2.2.1 Point spread function models

##### Scalar PSF Model

For optical systems with low numerical aperture (NA), the intensity profile in the focal plane and the complex signal in the pupil plane are related via the Fourier transform, as described as the Fresnel approximation[29]. A necessary condition for the validity of the Fresnel approximation is that the effect of light polarization on the diffracted images is negligible. The scalar PSF model can be related to the wavefront aberration *ϕ(*x*)* as

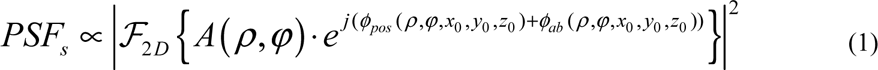

Where *ℱ_2D_* is the 2D Fourier transform, 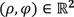 is the normalized polar coordinate in the back focal plane (BFP). *(*x*_O_*, **y*_O_*, **z*_O_)* is the 3D position of the point source in the sample plane. *PSF_s_* is the intensity of the optical field in the focal plane, *A* is the amplitude of the pupil plane. *ϕ_pos_(ρ*, *φ*, **x*_O_*, **y*_O_*, **z*_O_)* indicates the phase shift correlated with the emitter position and objective

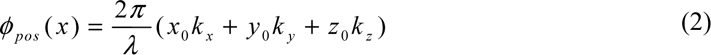

With

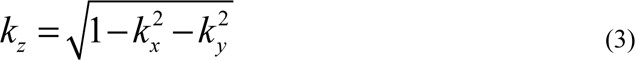

Where *λ* is the wavelength, and (*k_x_*, *k_y_*) are the **x** and **y** components of the unit wave vector. *ϕ_ab_*(*x*) represents the field-dependent aberrations and is expressed as a liner sum of variance normalized Zernike polynomials:

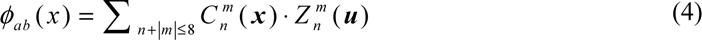

Where *x* = (**x*_O_*, **y*_O_*, **z*_O_*), ***x*** = (*x_o_*, *y_o_*, *z_o_*), 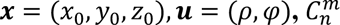 is the aberration coefficients, 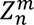 is the normalized Zernike polynomial, *n* is the radial order and *m* is the angular frequency.

##### Vectorial PSF Model

The scalar PSF model only focuses on the Fourier transform relationship between pupil and PSF according to the Fresnel approximation, while ignores the vector nature of light, which is applicable in low-NA situations. However, for high-NA optical systems[33] (e.g., NA ≥ 0.6), the vector nature of light cannot be neglected because the bending of the rays created by a lens introduces a significant **z** component of the electromagnetic field in the region behind the lens. We can model the PSF according to the vector theory of diffraction[34] by considering the **x**, **y**, **z** components of the field right after the lens separately for polarization vector with **x** and **y** components. We consider that each volume element of the sample represents an ensemble of independently emitting fluorophore molecules, which we described as uncorrelated dipole radiators (with random orientations) which can rotate or wobble freely. The PSF are treated as the summed image of three orthogonal dipoles: **x**, **y**, **z** directions. Thus, our calculation of the vectorially PSF simply consists of calculating and adding six intensity PSFs incoherently which is modulated by six field vectorial coefficients:

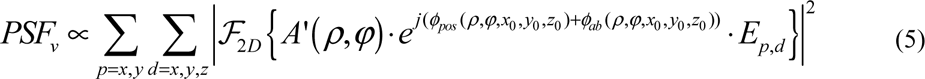

With

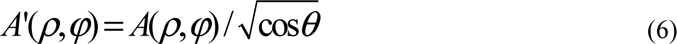

Where *A*’(*ρ*, *φ*) indicates the amplitude function with obliquity factor to account for the angle*θ* between the Poynting vector and the normal to the imaging plane. And the six coefficients *E_p_*_,*d*_ represent the polarization vectors of electrical field with components *p* = **x**, **y** in the image plane with orthogonal dipole components *d* = **x**, **y**, **z** contributes to components *p*:

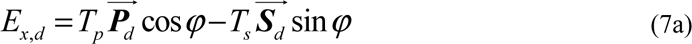

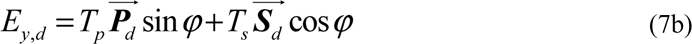

With

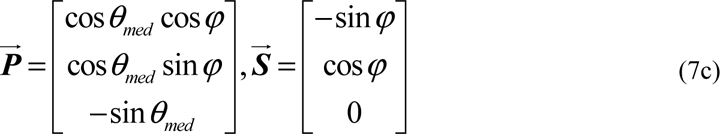

Here 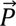 and 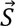 are the light field polarization vectors. *T_p_* and *T_s_* are the Fresnel transmission coefficients for *p* and *s* polarization. The six field components can be used to calculate the total electrical field energy at any point after the lens; in particular, they determine the intensity seen by an imaging plane in any orientation.

#### 2.2.2 MLE phase retrieval algorithm

MLE is an effective method to fit Poisson distribution models. We know that the measurement noise of the camera along pixels is independent and identically distributed with Poisson function, so the PSF intensity profile on the camera is also Poisson distributed, which can be accurately fitted by MLE especially for low signal-to-noise conditions.

The proper procedure with MLE phase retrieval is to adjust fitting parameters to minimize the MLE for the Poisson distribution. We use the L-M algorithm[25] to minimize the maximum-likelihood based loss-function given by

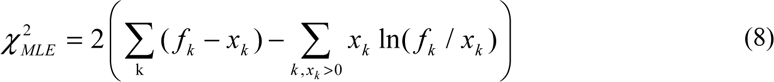

Where *f_k_* is the expected number of photons in pixel *k* from the model PSF function with parameters, and **x*_k_* is the measured number of photons in raw data. This function is minimized to find the maximum likelihood for the Poisson process.

### 2.3 Sample preparation

For 100nm-bead samples, we used the dark red fluorescent FluoSpheres beads (F8801) for fluorescent bead samples which were maximally excited at 589 nm. Dilute the beads in ultrapure water at a ratio of 1:10000, and vibrate in an ultrasonic cleaner for 5 mins. Next, 200 μl of the diluted beads were placed on a coverslip and allowed to stand for 10 mins. The absorbent tissue was then used to remove excess water and Prolong GlassDiamond Antifade (Thermo Fisher Scientific, Inc.) before sealing the coverslip.

### 2.4 Data acquisition

We tested the accuracy of VISPR from beads datasets. Before data collection, the deformable mirror (DM) was calibrated to introduce single-aberration modes with other random but non-primary Zernike coefficients to the imaging system based on Zernike polynomials. We utilized 21 Zernike modes within the Fringe order (amplitude ±1, units λ/2π), ranging from vertical astigmatism to third-order spherical aberration. After the fluorescent beads were distorted, data collection involved the following steps: 1) 20-30 bead stacks were collected for in-vitro phase retrieval (PR). Each bead stack was collected by moving the sample stage from −600 nm to 600 nm with a step size of 50 nm. 2) beads were captured for VISPR by moving the sample stage from −600 to 600 nm with a random step interval, i.e., the step sizes range from 2 to 7 nm randomly, to mimic in-situ SMLM raw data. 3) 50 repeated bead images for each depth from −600 to 600 nm with a 50 nm step were collected for validation. One frame per z position was collected at 20-35 ms exposure time.

## 3. Results

### 3.1 Simulation Verification

We first validate the feasibility of our VISPR extensively with single molecule simulations. Specifically, we generate a single-molecule randomly blinking dataset (Figure. 2a) based on the cubic-spline representation of a ground truth oblique astigmatic PSF model with a set of Zernike aberrations consisting of 21 Zernike modes (Wyant order[35]) and random axial range of ±800nm (4000 total photons/localization, 100 background photons per pixel, z-range −600 nm to 600 nm). Each molecule is added with a constant background and finally degraded by Poisson noise. In simulated experiments, VISPR demonstrates its proficiency by successfully estimating the ground truth pupil (Figure 2b) with a residual error of 13 nm (root-mean-square (RMS) error of the wavefront) and accurately determining the Zernike amplitude coefficients with an error of 11 nm for all 21 modes (Figure 2c). As shown in Figure S2, the VISPR PSF is close to reaching the Cramér-Rao lower bound (CRLB) under different aberration conditions.

**Figure 2.**
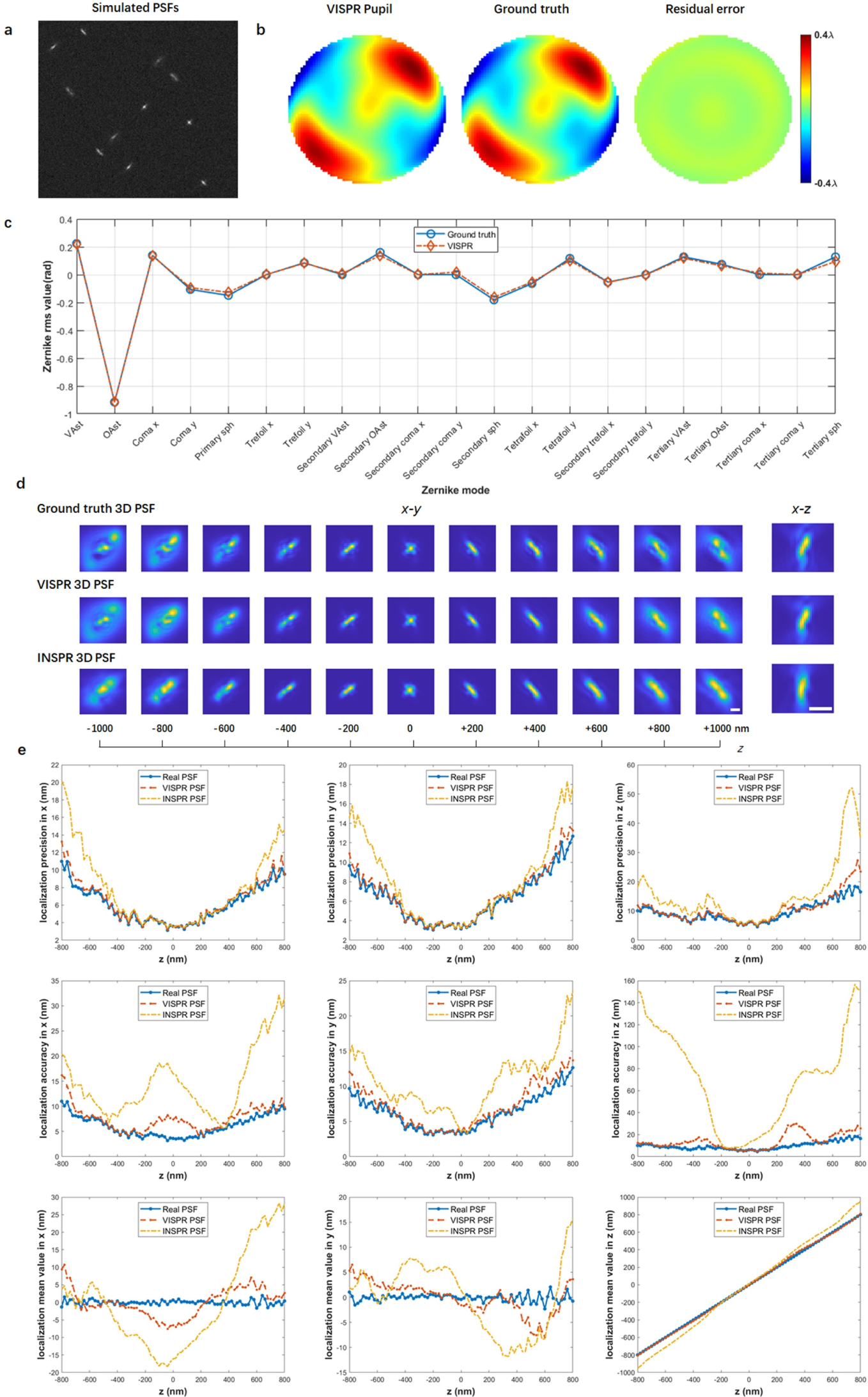
Performance quantification of VISPR on simulations. **a)** Simulated single-molecule dataset located randomly over an axial range from –800 to +800 nm with a known PSF model. **b)** Phase of the VISPR pupil (left), the ground truth pupil (middle) and the residual error (right). The RMSE is 11.33 nm. **c)** Coefficients of 21 Zernike modes retrieved by VISPR compared with the ground truth (red circles). **d)** x– y and x–z views of the Ground truth 3D PSF (top row), VISPR-retrieved 3D PSF (middle row), INSPR-retrieved PSF (bottom raw). Scale bar: 500nm. In the −400 to 400nm region, VISPR and INSPR are very similar to the real PSF. However, in the region beyond the z-axis 400nm, INSPR shows a severe deviation from the real PSF due to the lack of high-frequency information, while VISPR still maintains high similarity with the real PSF. **e)** Localization precision, accuracy and mean value in **x**, **y**, **z** positions at different axial positions with real PSF, VISPR and INSPR.

Next, we conduct a comparative analysis between the outcomes of VISPR and INSPR[28]. In the simulation experiment, within the −400 to +400 nm range, both INSPR and VISPR retrieved PSF demonstrate significant similarity with the ground truth PSF as shown in Figure 2d. As the depth along the z-axis extends beyond ±400 nm, the VISPR retrieved PSF still maintains a notably high degree of similarity to the real PSF due to taking into account the high-frequency information of the PSF. However, the INSPR PSF model introduces noticeable distortions with the ground truth PSF, due to inappropriate PSF model and failure to consider the high-frequency information of the data.

Moreover, we conduct a three-dimensional reconstruction of the simulated single-molecule datasets using separate ground truth 3D PSF, INSPR PSF, and VISPR PSF models. The evaluation encompasses localization precision, accuracy, and mean values in 3D at various axial positions, utilizing 100 simulated molecules at each axial position to comprehensively illustrate the performance of VISPR and INSPR as shown in Figure. 2e. The *localization precision* is calculated as the standard deviation of the difference between the fitted positions and ground truth positions. The *localization accuracy* is calculated as the root mean squared error between the fitted positions and ground truth positions. Figure. 2e shows that the evaluated **x**, **y**, **z** positions results localized by VISPR-retrieved 3D PSF model showed excellent similarity with the ground truth of the simulated data even in large z-axis. In contrast, substantial deviations are observed between the **x**, **y**, and **z** positions located by INSPR and the ground truth. Particularly noteworthy is that the mean z position of the INSPR PSF within the −400 to 400 nm range closely aligns with the real PSF; however, a considerable deviation becomes evident beyond the 400 nm range.

Subsequently, we further verify the performance of VISPR under conditions of low signal-background ratios considering the situation of conventional single-molecule experiments characterized by pronounced background noise. We test the Root-mean-square error (RMSE) between the VISPR fitted pupil and the ground truth pupil under varying signal-background ratio conditions (different photon numbers (I) and backgrounds (bg) conditions). As shown in Figure S3, even when confronted with a peak signal-to-background ratio (SBR) as low as 2:1 (2000 total photons, 100 background photons per pixel, z-range −600 nm to 600 nm), with a molecule count of 1000, the RMSE of the estimated pupil is also approximately 33nm. Furthermore, with an increase in input photons to more than 3000 molecules, the estimated pupil achieves higher accuracy, reaching around 20 nm. This analysis sheds light on the robustness and accuracy of VISPR under conditions reflective of real-world variations in signal-background ratios.

Furthermore, we utilize VISPR and INSPR to reconstruct the simulated microtubule structure respectively. The ground truth of the microtubules is obtained from the EPFL 2016 SMLM Challenge training dataset (http://bigwww.epfl.ch/smlm). Initially, we employ a ground truth oblique astigmatic PSF model to generate the simulated single-molecule dataset. Next, the single-molecule dataset of microtubules is inputted into VISPR and INSPR, and the 3D PSF model retrieved separately is employed to reconstruct the microtubule super-resolution image. The results reveal that within the −300 to 300nm region of the z-axis, the x-z reconstruction outcomes of VISPR (Figure. 3c and 3d) and INSPR (Figure. 3i and 3j) are closely aligned with the ground truth values. However, as the depth extends beyond 500 nm, the z-axis position of INSPR deviates significantly from the ground truth by approximately 200 nm (Figure 3k and 3l). In contrast, the results obtained with VISPR exhibit a good fit, closely aligning with the ground truth (Figure 3e and 3f). This observation underscores the effectiveness of VISPR in enabling the accurate blind super-resolution reconstruction of aberrated SMLM datasets.

**Figure 3.**
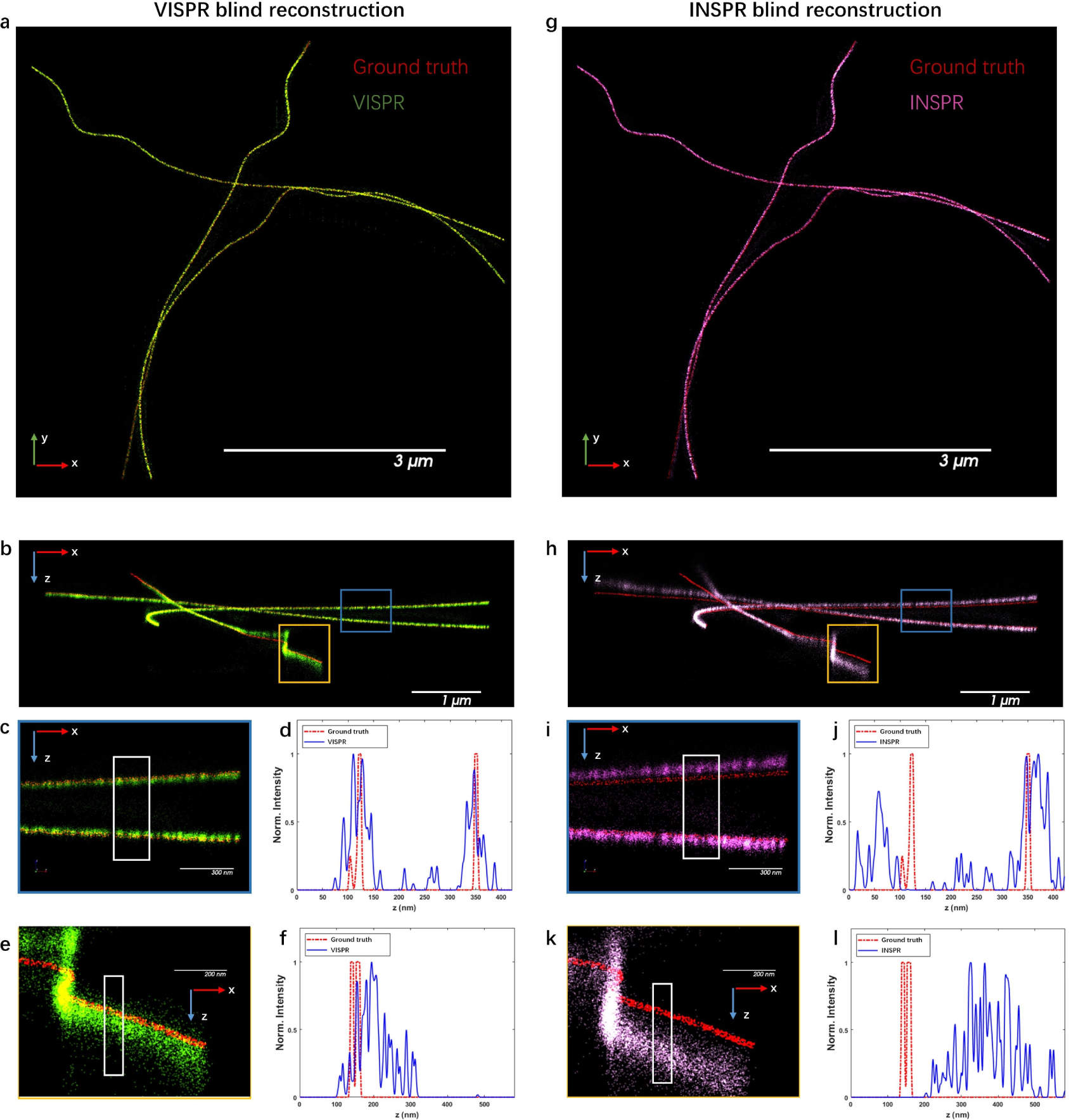
Comparison of the effectiveness of blind reconstruction of 3D SMLM microtubules between VISPR and INSPR. **a, b, g, h)** x-y and x-z overviews of the simulated microtubules resolved by VISPR and INSPR from the 3D SMLM data. **c, e)** Enlarged x-z views of the areas indicated by the blue and orange boxed regions in (b), respectively. **d, f)** Intensity profiles along the z directions within the white boxed regions in (c, e), comparing the VISPR resolved profiles (blue solid lines) with the ground truth (red dashed lines). **i, k)** Enlarged x-z views of the areas indicated by the blue and orange boxed regions in (h), respectively. **j, l)** Intensity profiles along the z directions within the white boxed regions in (i, k), comparing the INSPR resolved profiles (blue solid lines) with the ground truth (red dashed lines).

### 3.2 Experimental Verification

#### 3.2.1 Evaluation by Imaging Fluorescent Beads

To evaluate the performance of VISPR, experiments on 100 nm fluorescent nanospheres, which were settled on the upper surface of the cover slip, were conducted as a proof of concept. We added an oblique astigmatism-dominant aberration with other random but non-primary Zernike coefficients to the imaging system by deformable mirror (DM) (Figure. 4a). The validation experiment was divided into three parts: 1) We staged and captured the beads from −600 to 600 nm with a 50 nm step. The average of stacks of bead candidates with high photon intensity within the field of view (FOV) was utilized for in-vitro PSF/pupil retrieval using vectorial model and MLE fitting. This PR PSF and pupil were regarded as the ground truth of the system; 2) We staged and captured the beads from −600 to 600 nm with a random step interval, i.e., the step sizes range from 2 to 7 nm randomly, to mimic in-situ SMLM raw data (Figure. S4c). VISPR and INSPR methods were applied to get the in-situ PSF model and pupil function, respectively. It can be seen that in Figure 4a, both PR and VISPR can accurately retrieve the shape of the beads. In contrast, the shape of the INSPR PSF model at large depths (>300nm) deviates significantly from the PR PSF and real beads, because INSPR cannot retrieve the high-frequency information of the beads image; 3) We captured 50 repeated bead images for each depth from −600 to 600 nm with a 50 nm step. For each bead candidate in the FOV, PR PSF, VISPR PSF, and INSPR PSF models were utilized to fit the 3D coordinates at each depth, and the average values of **x**/**y**/**z** coordinates of the PR PSF result at each depth were regarded as the ground-truth coordinates for this specific bead candidate. Figure 4b shows that both PR and VISPR localization results exhibit a step-like upward trend, indicating the movement of the piezo positioner. Then, Localization precision (Δx/Δy/Δz) and corresponding accuracy (mean square error relative to the ground-truth coordinates) of PR/VISPR/INSPR fitting at each depth were calculated. For the VISPR/INSPR localization results, we added a global 3D displacement to the original fitting coordinates to compensate for the offset between the PR PSF and VISPR/INSPR PSF model. This is reasonable because the translation of the 3D reconstruction does not affect the final structural information. In Figure. 4c, we present a comparative analysis of the x-y-z axis localization precision and accuracy for beads using PR/VISPR/INSPR, respectively. Due to the sample stage’s random jitter, the z-position of the beads in the same z-axis area will also deviate, which will cause fluctuations in precision and accuracy. However, by comparing precision, accuracy and mean value at different z-axis positions, we still find that the average positions of VISPR and PR on the z-axis positioning can correctly reflect the movement of the stage. In contrast, the position of INSPR produces obvious deviations from the PR PSF, which illustrates the limitations of INSPR.

**Figure 4.**
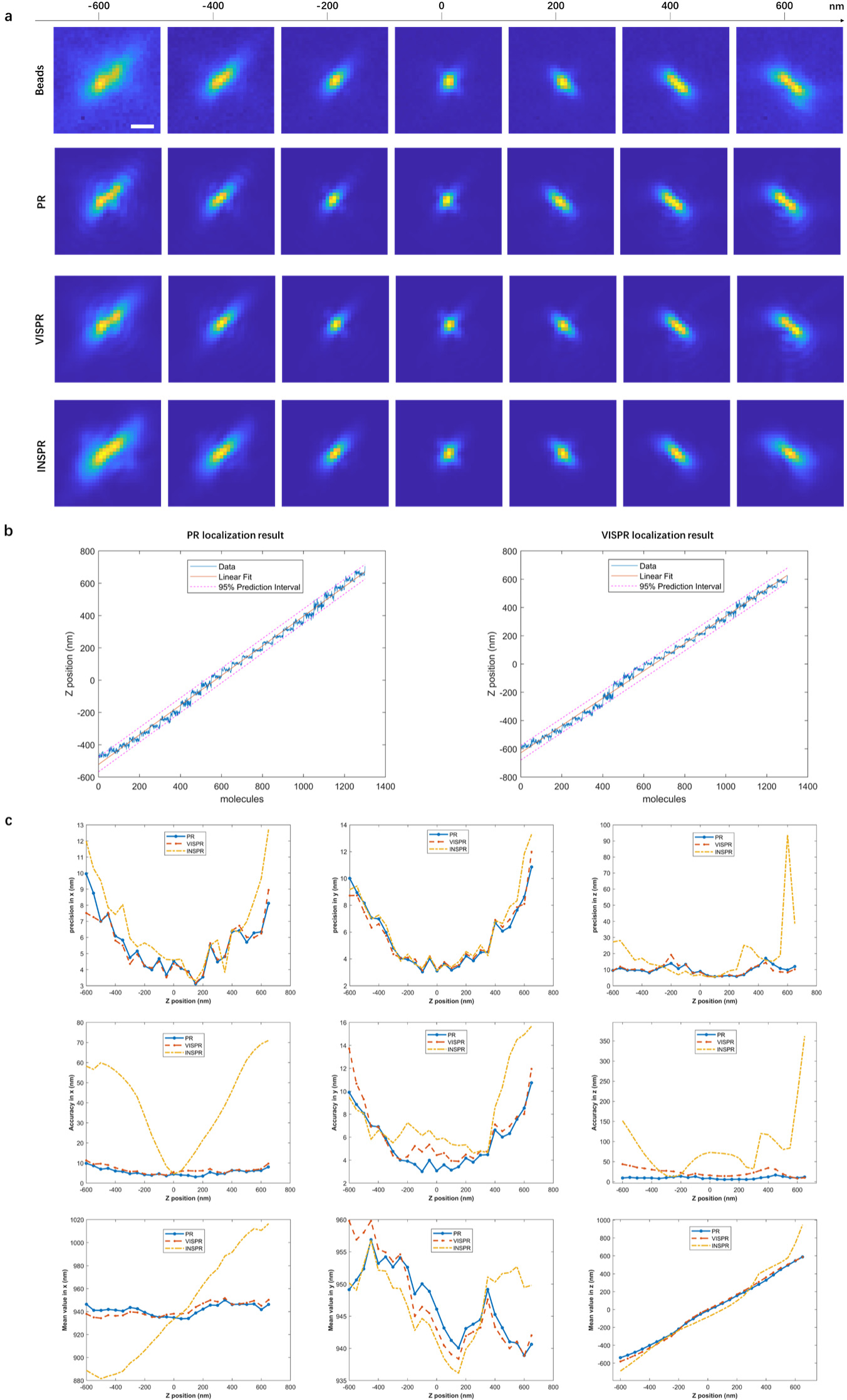
Experimental validation of VISPR on oblique astigmatism-dominant aberration. **a)** The original data of beads, the PSF obtained by PR, the PSF obtained by VISPR, and the PSF obtained by INSPR are presented at different z-axis depths. Scale bars: 500nm. **b)** The localizing results of the verification beads. For each depth from −600 to 600 nm with a 50 nm increment, 50 consecutive bead images are captured and reconstructed by PR, VISPR, and INSPR. Both PR and VISPR localization results exhibit a step-like upward trend, reflecting the movement of the piezo positioner. Linear fitting lines are applied to the positioning results, with all points falling within the 95% prediction area. **c)** Localization precision, accuracy and mean value in **x**, **y**, **z** positions at different axial positions with PR, VISPR and INSPR. This comprehensive analysis demonstrates the effectiveness and reliability of VISPR.

To further explore the robustness of VISPR, we conducted in-situ experiments with other types of aberrations. i.e., vertical astigmatism-dominant aberration with other random but non-primary Zernike coefficients as shown in Figure S4. The PSF estimated by VISPR demonstrated a clear linear relationship between the fitted z-position and the actual objective z-position. In contrast, INSPR exhibited distortion, indicating a potential misalignment of the PSF model with the actual data.

#### 3.2.2 Evaluation by Imaging Nup98 in U-2 OS cells

VISPR empowers the measurement and analysis of both system and sample-induced aberrations, facilitating the extraction of the authentic 3D PSF model in the situation of SMLM. This capability enables precise three-dimensional positioning of single molecules. To further validate the robustness and efficacy of VISPR, we conducted experiments focusing on imaging immunofluorescence-labeled nucleoporin Nup98 in U-2 OS cells.

We first reconstructed a high-resolution 3D volume of Nup98 at the bottom surface of the nucleus, ∼3 µm above the coverslip (Figure. 5a). Our observations revealed distinctive ring-like structures that spanned the lower surface of the nuclear envelope, featuring subtle invaginations and undulations (Figure. S5). Notably, as shown in Figure. 5c, due to the proximity of the Nuclear Pore Complex (NPC) to the coverslip, the bottom of the concave structure along the axial distribution displayed minimal disparities when reconstructed using both in-vitro phase retrieval (PR) methods and VISPR. However, increasing depth at the edge of the concave structure resulted in distorted and diffuse reconstruction with beads PR. In contrast, the reconstruction of VISPR exhibited consistent and converging axial distributions at the edge of the concave structure.

**Figure 5.**
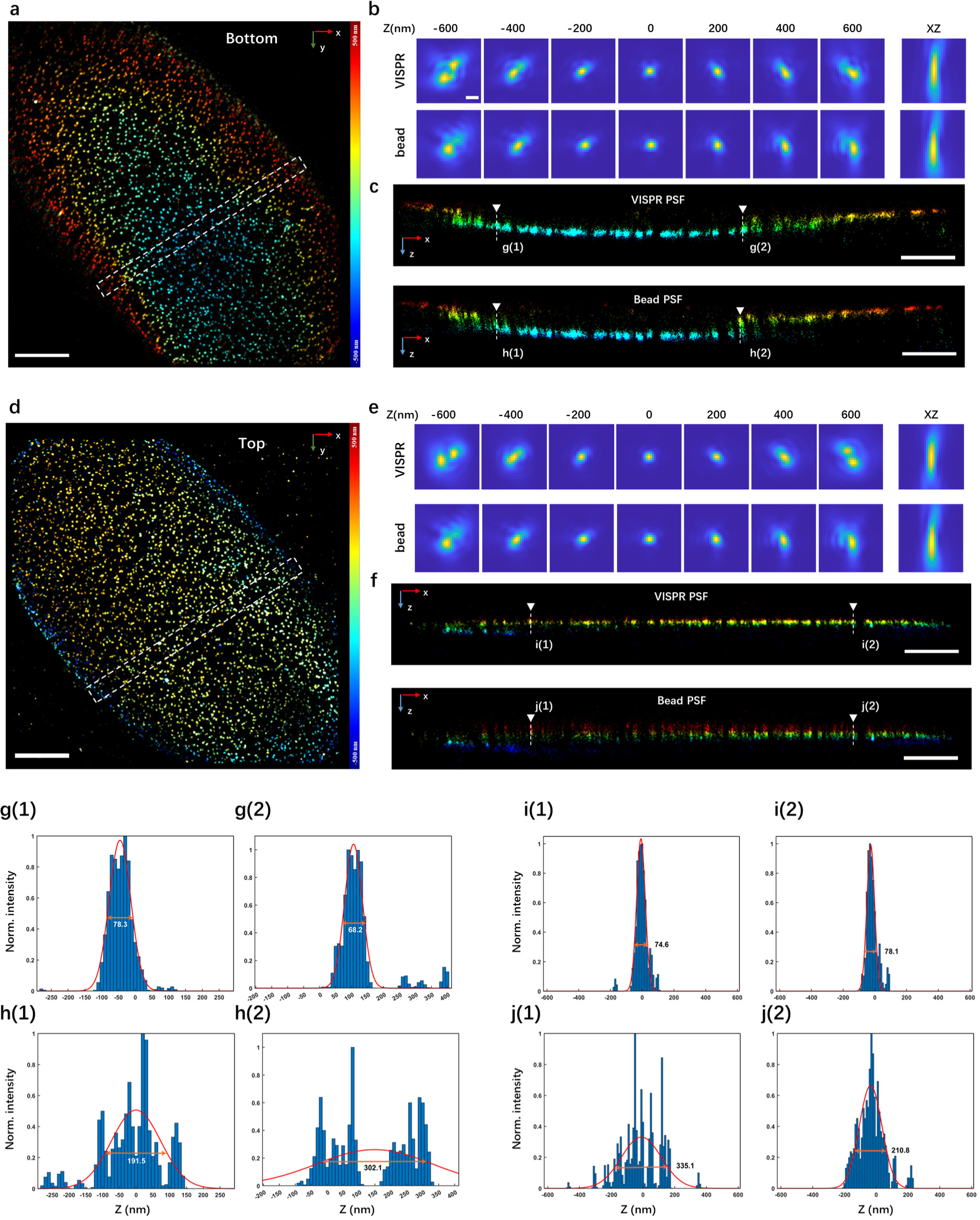
3D super-resolution reconstruction of immunofluorescence-labeled Nup98 on the nuclear envelope in U-2 OS cells using VISPR and the in vitro method. a, d), x–y overview of Nup98 at the bottom and top surface of the nucleus. b, e), the PSF of the bottom and top surface obtained by VISPR and the PSF obtained by beads are presented at different z-axis depths. c, f), x-z cross-section of the selected region in (a, d). g-j), Intensity profiles along the white dashed lines in c and f, which demonstrate the higher quality of VISPR PSF. Scale bars, 5 µm (a, d), 1 µm (c, f), 0.5 µm (b, e).

Subsequently, we proceeded to image Nup98 labeled with AF647 at the top region of the nucleus, ∼5 µm above the coverslip (Figure. 5d). This specific imaging condition, employing an oil immersion objective lens (NA=1.45) and standard imaging buffer (n=1.346), inherently introduced substantial sample-induced aberrations. We employed VISPR and the in-vitro phase-retrieval (PR) method based on fluorescent beads affixed to the coverslip to reconstruct the same field of view. Through comparative analysis, we found that the NPC reconstructed by VISPR exhibited a consistent distribution along the z-axis (Figure. 5f). In contrast, reconstructions using the in vitro PR approach showed a gradual spread and diffusion over a large depth range, accompanied by noticeable distortions and a reduction in resolution, as observed in the intensity profile (Figure. 5g-j).

By evaluating the reconstruction results on both the lower and upper surfaces of the nuclear membrane, VISPR demonstrates its capability to not only resolve the central structure of the nuclear pore within the horizontal axis but also maintain a consistent distribution along the z-axis. Despite the increasing influence of refractive index mismatch as depth increases, VISPR’s reconstructed structure exhibits remarkable consistency. These results underscore VISPR’s capacity in obtaining the accurate PSF model even at considerable depths. Conversely, the in-vitro method neglects the consideration of sample-induced aberrations such as refraction mismatch, leading to distortion and significant spreading of reconstructions along the extensive z-direction.

To further explain this difference, we compared the VISPR-retrieved PSF models with the in-vitro one, as illustrated in Figure 5b and e. Analyzing the VISPR PSF of both the bottom and top surfaces of the nucleus reveals that, with an increase in imaging depth, the extent of sample-induced aberrations, such as spherical and coma aberrations from optical sections, intensifies. The VISPR-retrieved pupils effectively captured this depth-dependent variation in sample-induced aberrations, their decomposed Zernike amplitudes, and the corresponding axially stretched vectorial PSFs. In contrast, the PSF retrieved from fluorescent beads only measured instrument imperfections, but failed to consider sample-induced aberrations and their depth-dependent variations such as refraction mismatch. Given that sample-induced aberrations can vary significantly from specimen to specimen, resulting in a degradation of axial reconstruction for in-vitro PR algorithms, our results underscore the consistent ability of VISPR to achieve accurate and high-resolution 3D reconstructions.

## 4. Conclusion

In this work, we propose VISPR, a robust in-situ phase retrieval framework that can directly retrieve 3D PSF models considering both system and sample-induced aberrations from raw single-molecule datasets. VISPR facilitates the accurate positioning of single molecules in three dimensions, thereby enabling accurate 3D SMLM localization. Our approach is validated through simulation datasets and practical experiments. In the simulation experiment, we confirm that VISPR exhibits superior fitting ability compared to INSPR, which is attributed to enhancements in both the model and phase retrieval algorithms. Importantly, VISPR ensures the absence of distortion in the recovery of 3D PSF models, even at significant depths, distinguishing it from INSPR. With the help of the vectorial PSF model and MLE phase retrieval algorithm, VISPR effectively addresses issues related to model deviation and the absence of high-frequency information at substantial depths along the z-axis. Furthermore, even in scenarios with low signal-to-noise ratios, VISPR demonstrates robust performance. It is essential to note that VISPR’s efficacy relies on single-molecule data within a specific z-axis range, and potential limitations may arise if the majority of single-molecule data cluster around the same z-axis position.

Traditional phase retrieval methods relying on beads are inadequate for SMLM data at large depths such as intra- and extracellular cellular targets deep inside tissues. This inadequacy stems from the fact that many biological samples exhibit significant variations in refractive index, a feature not captured by beads. VISPR, in contrast, emerges as a robust solution capable of accurately retrieving the in-situ 3D PSF model. Its strength lies in its simultaneous consideration of system and sample-induced aberrations. Additionally, VISPR improves the accuracy of the PSF model by solving the problem of PSF model distortion at large depths and considering the high-frequency information of the dataset. Noteworthy is VISPR’s versatility, demonstrating applicability in low signal-to-noise ratio SMLM situations and compatibility with various microscope modalities. This versatility provides invaluable insights for optimizing optical system performance, correcting and analyzing aberrations induced by the system and samples, estimating single-molecule properties, and enhancing imaging contrast and resolution.

## Code availability

Data underlying the results presented in this paper are not publicly available at this time but may be obtained from the authors upon reasonable request.

## Author contributions

X.Y., H.Z. and Y.S. conceived and initiated the project. X.Y., H.Z. and Y.S. designed and built the optical system. X.Y. wrote the software control code. X.Y., H.Z., Y.S. and H.W. designed and performed simulations and experiments. X.Y., H.Z. analyzed the data. X.Y., Y.S. and H.W prepared the experimental samples. X.Y., and H.Z., wrote the article. C.K. and X.L. guided and supervised the whole project. All authors wrote the paper and contributed to the scientific discussion and revision of the article.

