## Supplemental information for "Accurate 3D SMLM localization via Vectorial In-situ PSF Retrieval and Aberration Assessment"

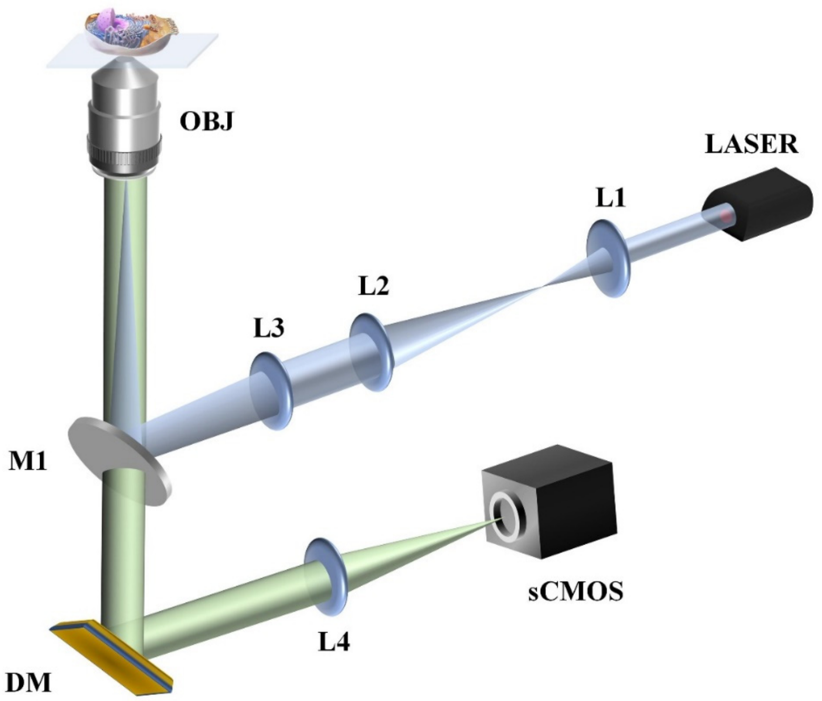


**Figure S1. Setup diagram.** L1–L3: lenses in the excitation path; L4: lenses in the emission path; M1: dichroic mirrors; OBJ: objective lens (Olympus, UPLXAPO100XO); DM: deformable mirror; sCMOS: scientific CMOS (Hamamatsu, ORCA-FusionBT C15440-20UP)

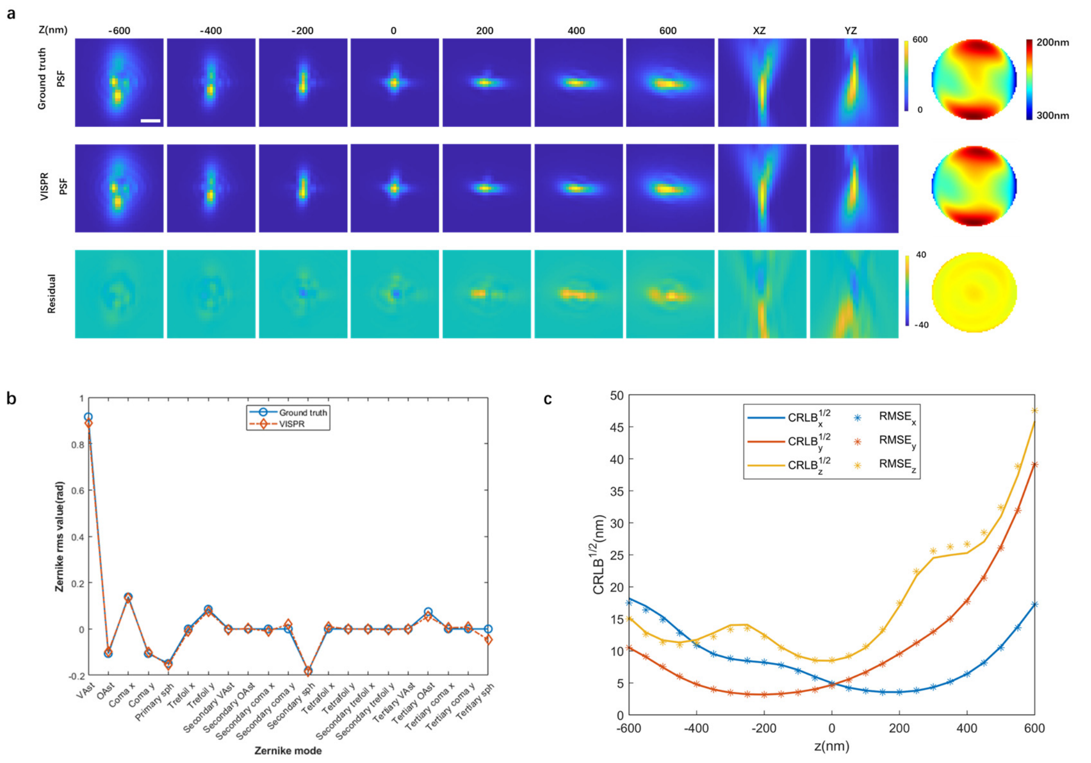


**Figure S2. Performance of VISPR estimated from simulated single-molecule blinking data. a)** Comparison of the ground truth PSF and VISPR estimated PSF from single molecules located in the axial range from -600nm to +600nm. The simulated data is randomly distributed from -600 to 600nm on the z-axis with a total of 4000 photons and a background level of 100. Scale bar: 500nm. **b)** Comparison of the ground truth coefficients of the 21 Zernike modes (blue line) and the amplitudes of the 21 Zernike modes retrieved from VISPR (orange line). **c)** Localization precision of 3D positions. 1000 repeated fitting calculations were performed to determine the localization precision. CRLB is the Cramér-Rao lower bound.

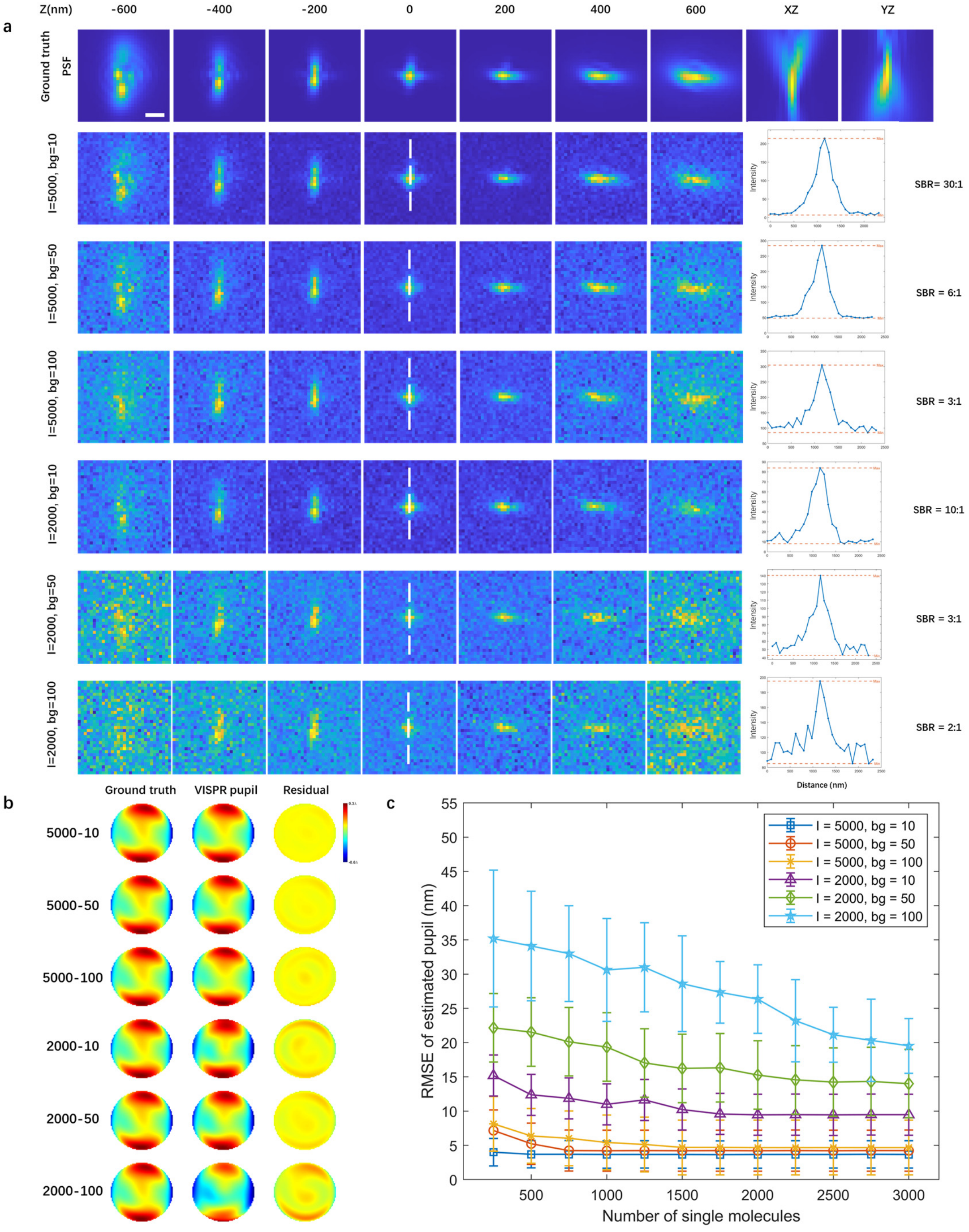


**Figure S3. Performance of VISPR under varying signal-background ratios. a)** Display of the ground truth PSF in different photon (I) and background (bg) conditions. In each condition, the intensity profile along the white dashed lines at the 0nm position PSF is analyzed to calculate the ratio of the peak signal to background. Notably, under the condition of 200 photons and 100 background, the peak signal-to-background ratio (SBR) plummets to as low as 2:1. Scale bar: 500nm. **b)** Comparison of the ground truth pupil and VISPR estimated pupil from single molecules located in the axial range from -600nm to +600nm under diverse signal-background ratios. **c)** Root-mean-square error (RMSE) between the VISPR estimated pupil and the ground truth pupil in different photon (I) and background (bg) conditions. In each condition, the ground truth pupil is randomly sampled from different combinations of 21 Zernike amplitudes for each trial, amounting to 11 trials in total. This comprehensive analysis sheds light on the robustness and accuracy of VISPR under conditions reflective of real-world variations in signal-background ratios.

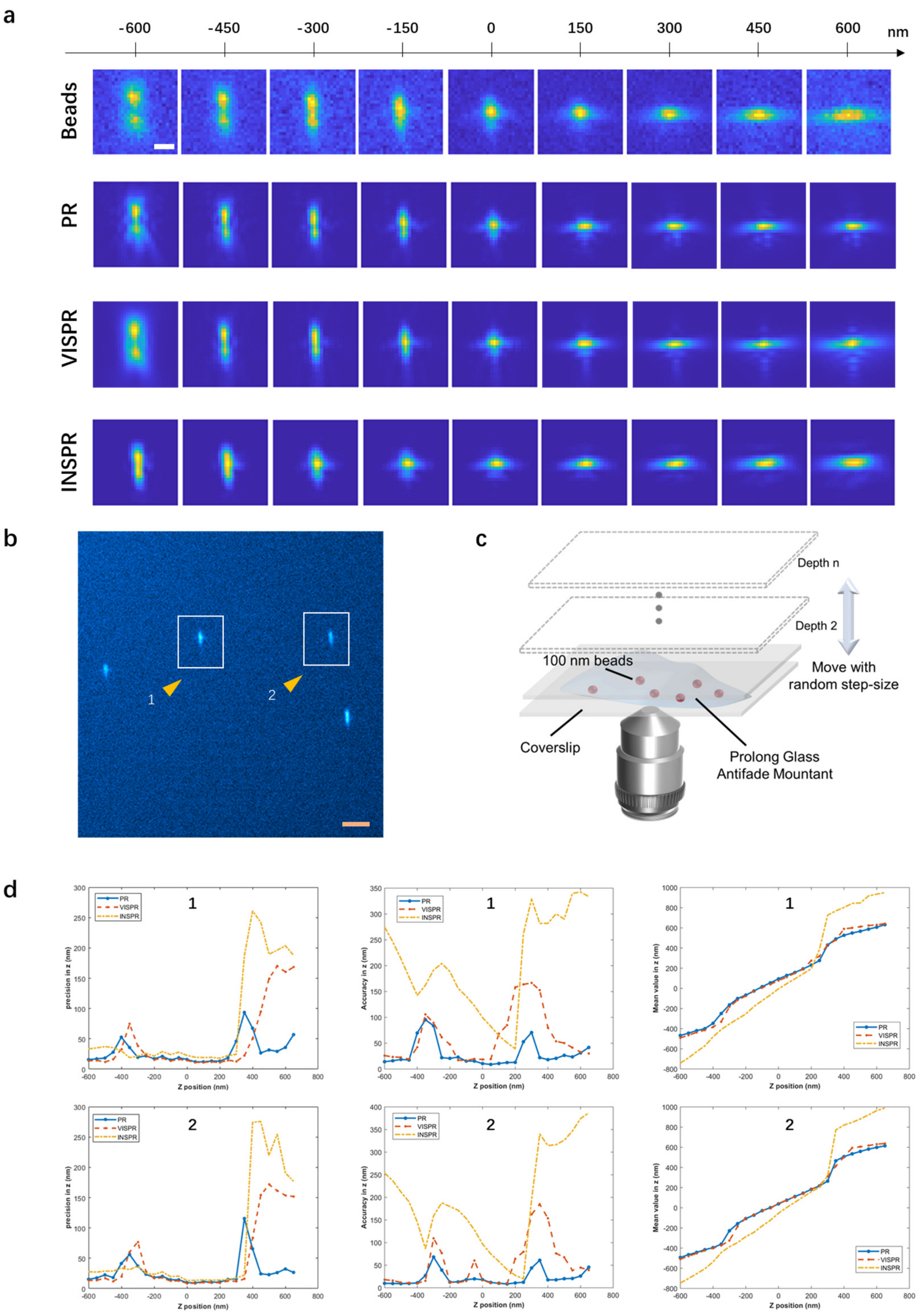


**Figure S4. Experimental validation of VISPR on vertical astigmatism-dominant aberration.**

**a)** The original data of beads, the PSF obtained by PR, the PSF obtained by VISPR, and the PSF obtained by INSPR are presented at different z-axis depths. **b)** Partial FOV showing the sample with sparse single beads. The localization precision, accuracy and mean value of the beads pointed by the yellow arrows are measured. **c)** A sketch of the sample used to measure localization precision at different depth. The 100 nm particles are on the upper surface of the cover slip and moved by a Piezo stage to different axial depths. **d)** Localization precision, accuracy and mean value curve across the depth of field of the two candidate molecules. Scale bars: 2 μm for b and 500 nm for others.

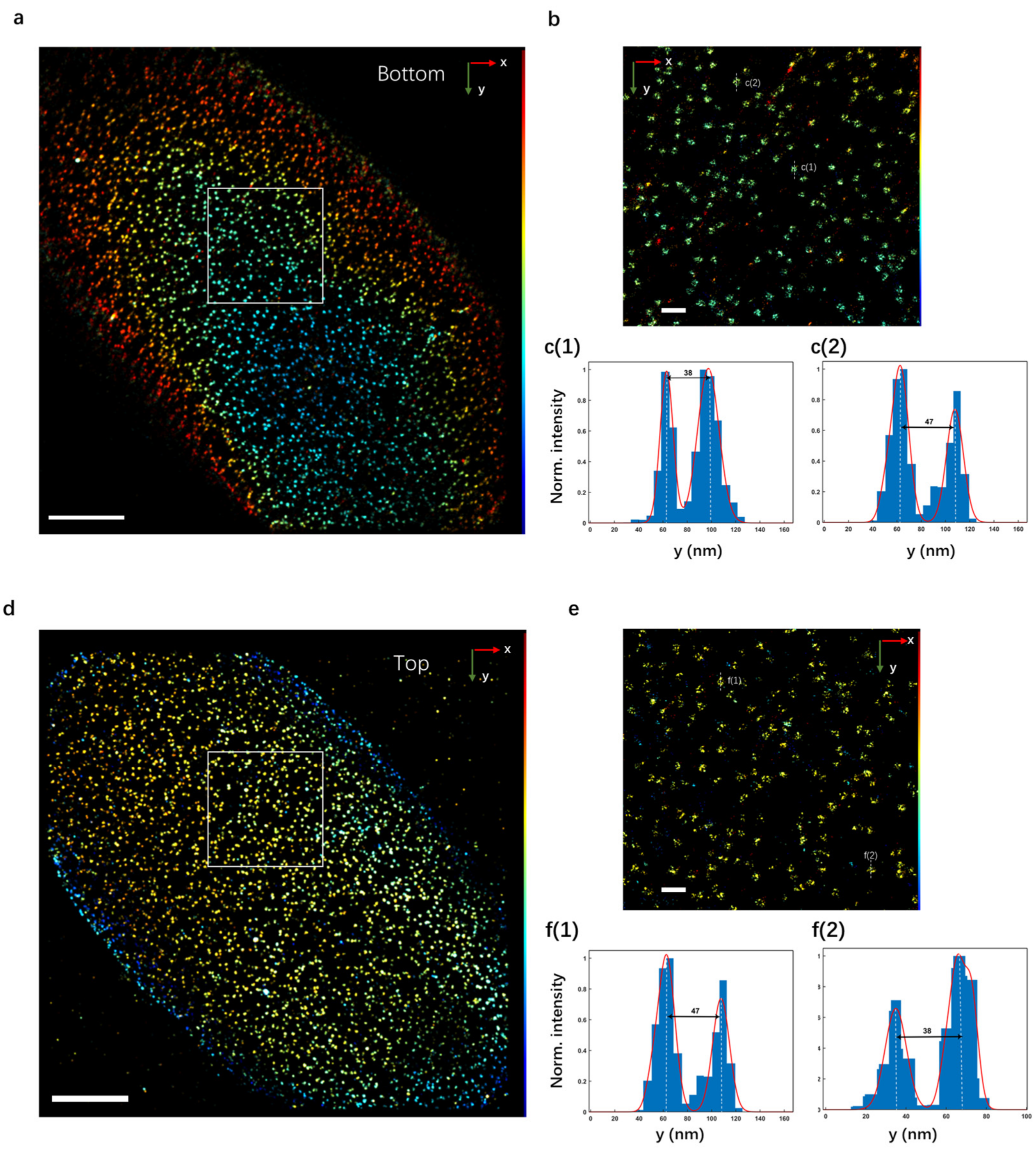


**Figure S5. 3D super-resolution reconstruction of immunofluorescence-labeled Nup98 on the nuclear envelope in U-2 OS cells using VISPR. a, d),** x–y overview of Nup98 at the bottom and top surface of the nucleus. **b, e),** Subregions, as indicated by the white boxed regions in a and d, showing the hollow structure of Nup98**,** showing distinctive ring-like structures. **c, f),** Intensity profile along the white dashed line in b and e, featuring subtle invaginations and undulations.
